# Single-cell RNA-seq highlights heterogeneity in human primary Wharton’s Jelly mesenchymal stem/stromal cells cultured *in vitro*

**DOI:** 10.1101/723130

**Authors:** Changbin Sun, Lei Wang, Hailun Wang, Tingrun Huang, Xi Zhang

## Abstract

Mesenchymal Stem/Stromal cells (MSCs) are multipotent cells with promising application potential in regenerative medicine and immunomodulation. However, MSCs cultured *in vitro* exhibit functional heterogeneity. The underlying molecular mechanisms that define MSC heterogeneity remain unclear. Here, we investigated gene-expression heterogeneity of human primary Wharton’s Jelly-derived MSCs (WJMSCs) cultured *in vitro* via single-cell RNA-seq. At the single-cell level, highly variable genes (HVGs) are associated with functional characteristics of classic MSCs. Differentially expressed genes analysis revealed the existence of several distinct subpopulations exhibit different functional characteristics associated with proliferation, development, and inflammation response. By comparing our WJMSCs data with a public available adipose-derived MSCs (ADMSCs) single cell transcriptomic data, we found that HVGs from these two studies are largely overlapped and have similar functional enrichment. Taken together, these results suggested that these HVGs hold the potential to be used as candidate markers for further potency association studies.

## INTRODUCTION

Mesenchymal Stem/Stromal cells (MSCs) are multipotent with self-renewal capacity and can be derived from various tissues, including bone marrow (Prockop, 1997), adipose tissue (Zuk et al., 2002), umbilical cord (Romanov et al., 2003; Wang et al., 2004), placenta (Fukuchi et al., 2004) etc. Besides the multi-lineage potential to differentiate into various cell types, such as chondrocytes, osteocytes, adipocytes, myocytes, and neuronal cells (Gao et al., 2016; Pittenger et al., 1999), MSCs could modulate immune cell response via interaction with lymphocytes from both the innate and adaptive immune system to deliver immunosuppressive and anti-inflammatory effects after homing to sites of inflammation *in vivo* (Aggarwal and Pittenger, 2005; Ren et al., 2008). Furthermore, human MSCs could be cultured in large scale and have minimal functional loss after long-term cryopreservation (Ikebe and Suzuki, 2014; Parekkadan and Milwid, 2010). Therefore, MSCs demonstrate promising utilization potential and are ideal cell types in both fundamental and translational biology fields, such as developmental biology, cellular therapy, immunomodulation, and regenerative medicine (Abdallah and Kassem, 2008; Baksh et al., 2004). Currently, more than 700 clinical trials have been registered in ClinicalTrials.gov (http://www.clinicaltrials.gov), which utilize MSCs for cellular therapy. Transplantation of MSCs demonstrates no obvious adverse effect, regardless of allogeneic or autologous cell origin, and has been extensively explored in treatment of various disease types, such as bone and cartilage defects (Krampera et al., 2006; Wakitani et al., 2002), cardiovascular disease (Chen et al., 2004; Ranganath et al., 2012), neurological degeneration (Karussis et al., 2010; Mazzini et al., 2010), liver disorder (Kharaziha et al., 2009), and immunological diseases (Ghannam et al., 2010; Le Blanc et al., 2008) with encouraging clinical outcomes. Several MSC-based products have been approved or conditionally approved in certain country or district to treat disorders, such as graft versus host disease (GvHD), Crohn’s-related enterocutaneous fistular disease (Galipeau and Sensebe, 2018).

The minimal criteria for defining multipotent MSCs was published by International Society for Cellular Therapy (ISCT) in 2006 (Dominici et al., 2006), which is widely accepted and adopted in both basic research and industrial application. However, it only defines basic morphological and functional characteristic. More and more research works have recognized that MSCs populations exhibit tissue-to-tissue functional variation (Jin et al., 2013; Yoo et al., 2009), as well as inter-population heterogeneity when using current markers to define MSCs, which makes it difficult to predict cell population dynamics and functional alterations after extended culture or exposed to extrinsic factors (Russell et al., 2010; Samsonraj et al., 2015). Functional heterogeneity coupled with large-scale expansion in clinical manufacturing process may explain, in part, why data across MSC-based clinical trials are largely incongruent (Phinney, 2012). During MSCs culturing and passaging, the competition and balance between different subpopulations may change, resulting in decrease in proportion or even loss of certain subpopulations and ultimately leading to alterations in cell function and treatment outcomes in clinical studies (Bustos et al., 2014; Sethe et al., 2006). Therefore, there is an urgent need to elucidate whether certain MSC subtype, or a cocktail of defined population of different subtypes can demonstrate effectiveness during cellular therapy and tissue engineering. Investigation into the underlying molecular mechanisms that define MSCs heterogeneity will facilitate subtypes identification and improve methods for cell isolation and expansion *in vitro*. By in-depth analysis of cell quality attributes, it will also help to interpret the results from clinical trials and eventually improve clinical efficacy of MSCs products.

Recently, single-cell RNA sequencing (scRNA-seq) technology, which allows massive parallel analysis of gene expression profiles at single-cell level, has become a powerful tool in investigating tissue and cell heterogeneity. It provides unprecedented opportunities for identifying subpopulations that share a common gene-expression profile in a heterogeneous cell population (Tang et al., 2009). Here, we investigated the gene-expression profile via scRNA-seq of human primary WJMSCs cultured in vitro from three donors. In contrast to other sources derived MSCs, WJMSCs, which isolated from previously discarded umbilical cord tissues, bear higher proliferation rate and the strongest immunomodulatory effect, making them an attractive alternative source of MSCs for clinical research and application (Li et al., 2014; Troyer and Weiss, 2008). Meanwhile, we analyzed public available ADMSC scRNA-seq data (Liu et al., 2019), and performed transcriptome comparison between WJMSCs and ADMSCs at the single-cell level. GO enrichment analysis of highly variable genes (HVGs) obtained from WJMSCs revealed that those gene products are significantly enriched in extracellular region with binding function, involved in developmental process, signal transduction, cell proliferation, etc. biological processes. Moreover, pathway analysis showed that those HVGs are associated with functional characteristics of classic MSCs, such as inflammation mediated by chemokine and cytokine signaling, integrin signaling, and angiogenesis. After regressing out the batch and cell cycle effects, those HVGs were used for dimension reduction and clustering analysis to identify candidate subpopulations. Differentially expressed genes analysis revealed the existence of several distinct subpopulations of MSCs that exhibit diverse functional characteristics related to proliferation, development, and inflammation response etc. Although WJMSCs and ADMSCs were derived from different tissues and displaying different differentiation potency, their HVGs were largely overlapped and had similar functional enrichment. Taken together, these results indicated that genes expression are highly varied among individual MSCs in culture. They are involved in different signaling pathways regulating individual cells response to the extracellular environment, which could eventually impact on population differentiation behavior and immunomodulation potency. Thus, those genes may serve as candidate markers for further potency association study.

## RESULTS

### Overview of WJMSCs Single-cell RNA Sequencing Data

To investigate into inter-population heterogeneity in primary cultured WJMSCs at the single-cell transcriptome level, primary cells isolated from three human umbilical cord (two females and one male, named as UC1, UC2, UC3 respectively) were collected and used for scRNA-seq. Total about 5 × 10^8^ raw reads with high quality for each donor were obtained (Figure S1A). Mapping these reads to human GRCh38 genome, average about 56.90 % and 61.03 % reads were mapped confidently to transcriptome and exonic regions, respectively (Figures S1B and S1C). Briefly, total 6 878 cells (filtered matrix) were obtained from the three donors, average 2 293 cells for each, with 209 769 mean reads, 38 983 medium unique molecular identifiers (UMI) counts and 6 279 median genes per cell (Figures S1D-S1F), suggesting that our data were of high quality.

According to the minimal criteria proposed by the International Society for Cellular Therapy (ISCT) in 2006 (Dominici et al., 2006), MSC must express three positive markers, i.e. *CD105* (*ENG*), *CD73* (*NT5E*) and *CD90* (*THY1*), and lack expression of several negative genes, including CD45 (*PTPRC*), *CD34*, *CD14* or *CD11b* (*ITGAM)*, *CD79a* (*CD79A*) or *CD19* and *HLA-DR* (*HLA-DRA* and *HLA-DRB1*). When we looked at the expression of those markers in our raw data, we saw the expression of those positive markers (UMI > 0), while negative genes were not expressed (UMI = 0) in most cells (Figure 1A). Next, we ranked cluster of differentiation (CD) genes by average normalized expression or percentage of cells with at least one UMI across all cells (Table S1). Classic cell surface markers for MSC definition, including *ENG*, *NT5E*, and *THY1*, as expected, belong to the top50 highly expressed CDs (Figure 1B). Among the CDs, integrins, such as *ITGB1*, *ITGA1*, *ITGA2*, *ITGA5*, etc., which play important role in MSCs morphology, migration, proliferation, differentiation and survival (Docheva et al., 2007; Hamidouche et al., 2009; Olivares-Navarrete et al., 2011), also highly expressed in WJMSC population (Figure 1B and Table S1). In addition, we assayed the tri-lineage capability of the cultured WJMSCs for scRNA-seq, and results confirmed that they have the potency to differentiate into osteoblasts, adipocytes and chondroblasts *in vitro* (Figure S1G).

**Figure 1.**
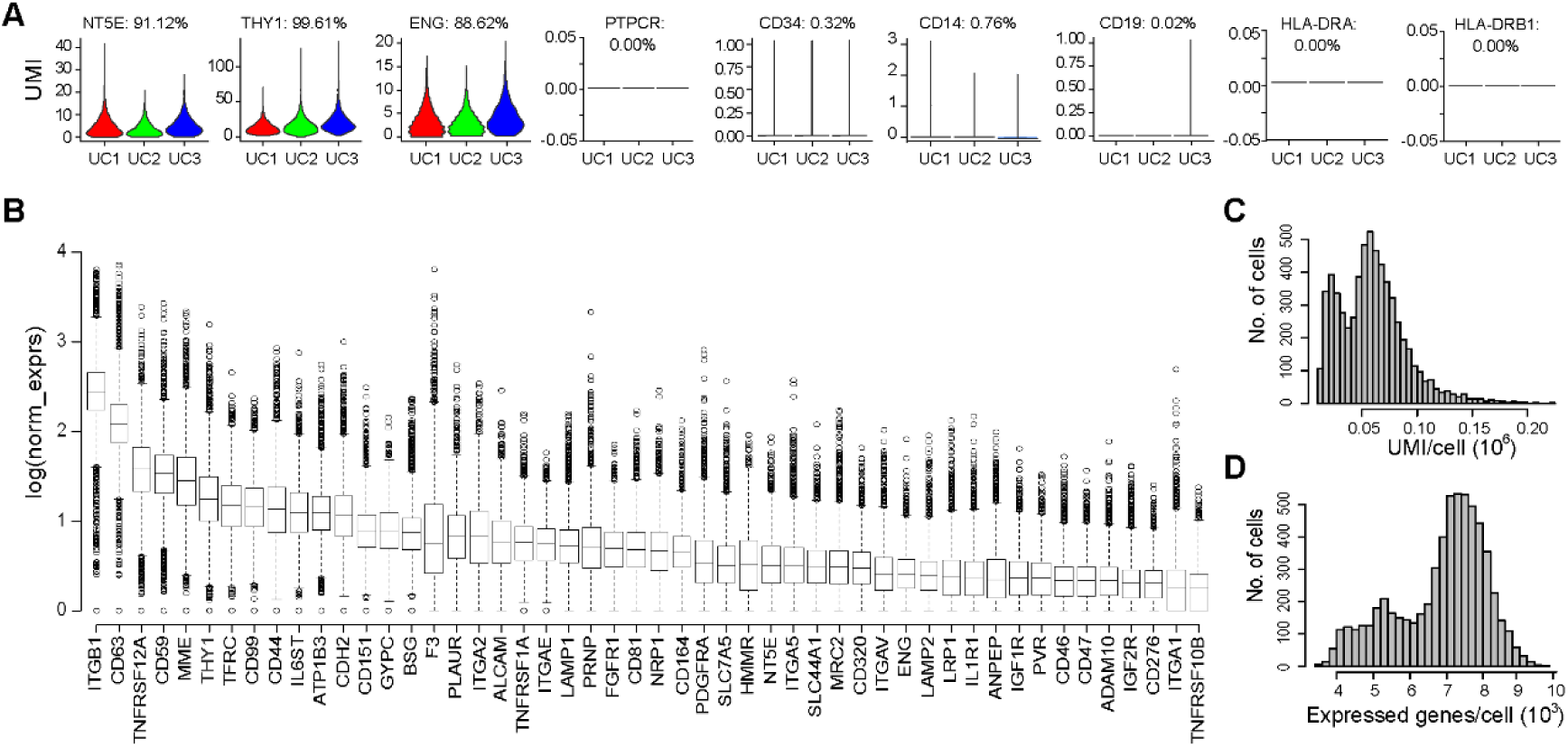
Overview of WJMSC single-cell RNA-seq data. (A) Expression of marker genes in the three samples. Number on the top showing percentage of cells with at least one UMI. (B) Boxplot showing top 50 cluster of differentiation (CD) genes ranked by average normalized expression. (C) Distribution of UMI cross cells after pre-processing to filter out low-quality cells. (D) Distribution of expressed genes after pre-processing to filter out low-abundance genes with mean-based method (genes with means more than 0.1 were retained).

For further analysis, we filtered outlier cells using the median absolute deviation from the median total library size (logarithmic scale), total gene numbers (logarithmic scale), as well as mitochondrial percentage for each donor (Lun et al., 2016). Totally, 702 outlier cells were removed and 6 176 single cells were kept by median absolute deviation method. Considering none or low abundant expressed genes across cells, we also integrated these three data together and removed any gene with average expression less than 0.1 UMI. Finally, 6 176 high quality single cells with 11 458 expressed genes were passed on to downstream analysis. Across the cells, number of UMI per cell ranged from 13 121 to 221 432, and number of genes from 3 543 to 9 775 (Figures 1C and 1D).

### Highly Variable Genes Identified in WJMSCs

Considering cell cycle effect may influence gene expression, we first assigned cell cycle phases state to each cell. Results showed that average 22.98 %, 34.51 %, and 42.51 % cells assigned to G1, G2/M, and S cell cycle phase, respectively (Figure 2A), suggesting that i*n vitro* cultured WJMSCs are highly proliferated population. Principle components (PCs) analyzed without removing unwanted sources of variation demonstrated that PC1, counting for 23.86 % variance, is mainly caused by cell cycle effect (Figure 2B), while PC2 counting for 10.10 % variance (Figure 2C), results from donor-to-donor variation or batch effect. Thus, we selected overlapped highly variable genes among each phase for each donor as mentioned in the supplemental method section and totally got 770 genes defined as HVGs for the following analysis (Figures S2A-S2D and Table S3).

**Figure 2.**
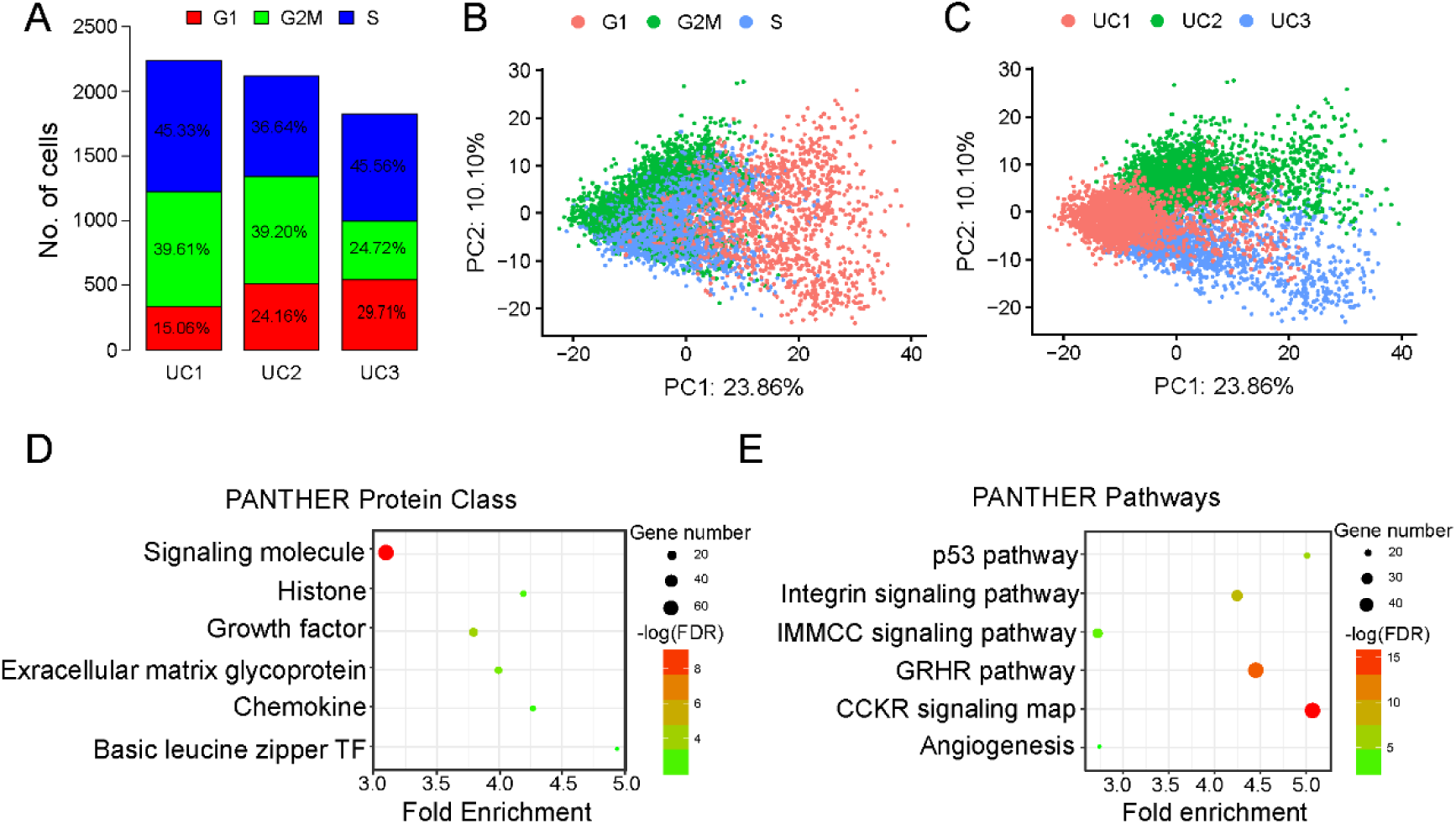
Heterogeneity and highly variable genes in WJMSCs. (A) Phases of cell cycle assigned for each of the three samples. (B and C) Cell cycle effects (B) and batch effects (C) represent the dominant source of heterogeneity in primary cultured WJ-MSCs population. (D)Results of pathway enrichment analysis for highly variable genes identified in WJMSCs. (E) Results of protein class enrichment analysis for highly variable genes identified in WJMSCs. IMMC: Inflammation mediated by chemokine and cytokine; GRHR: Gonadotropin-releasing hormone receptor.

Highly variable genes (HVGs) exhibiting high variability across cells represent heterogeneous features within a cell population (Pijuan Sala et al., 2019; Yip et al., 2018). Here, we investigated gene functional enrichment of HVGs identified in WJMSCs population. Interestingly, protein class analysis demonstrated that those genes were overrepresented in signaling molecules, growth factors, extracellular matrix protein, chemokine, histone, and basic leucine zipper transcription factor (Figure 2D). Besides, pathway analysis exhibited that these highly variable genes expressed cross cells were enriched in integrin signaling pathway, inflammation mediated by chemokine and cytokine signaling pathway, gonadotropin-releasing hormone receptor pathway, p53 pathway and angiogenesis (Figure 2E). Furthermore, GO enrichment analysis showed that those HVGs are significantly enriched in extracellular region (Figure S2E) with binding function, such as protein binding, and cytokine receptor binding, etc. (Figure S2G), involved in biological processes like developmental process, signal transduction, cellular component morphogenesis, cell communication, cell proliferation, etc. (Figure S2F). Micro-environmental interaction is crucial for morphogenesis, cell differentiation, homeostasis, cell growth (Frantz et al., 2010; Rozario and DeSimone, 2010). Therefore, variations in the expression of those extracellular functioning genes identified in our analysis could influence interaction of MSCs with micro-environment and cell fate determination (Even-Ram et al., 2006; Guilak et al., 2009). Furthermore, our results showed that highly variable genes in WJMSC population were enriched in distinct biological functions of MSCs, such as anti-inflammation (Figure 2E), regeneration (Figure S2F), wound healing (Figure S2G), etc., which can potentially be separated and purified to test their therapeutic efficacy in clinical application.

### Characteristics of Candidate Subpopulations in WJMSCs

To remove batch and cell cycle effects, we scaled the data and performed linear regression to regress the effects out before candidate subpopulations clustering. Here, we used regularized negative binomial regression method to perform normalization and variance stabilization of our scRNA-seq data, which is an appropriate distribution to model UMI count data from a ‘homogeneous’ single cell population suggested by (Hafemeister and Satija, 2019). Results of nonlinear dimensional reduction performed by UAMP showed that the cells were obviously separated by cell cycle and batch effects before regression, while cells were well mixed after regression and scaling (Figure S3A), implying that those unwanted sources of variation have been effectively removed.

Next, we performed cell cluster analysis by a graph-based clustering approach (Macosko et al., 2015), and six candidate clusters in primary cultured WJMSCs were identified (Figure 3A). To study the molecular and functional characteristics of these candidate subpopulations in WJMSCs, we performed differentially expressed genes (DEGs) analysis among the six clusters (C0-C5). (Table S4, Figure 3B). Intriguingly, *MKI67* (Marker of Proliferation Ki-67), a gene strongly associated with cell proliferation and growth, expressed at higher levels in subpopulations C0 and C1 compared with others, implying that subpopulations C0 and C1 possess a higher proliferative capacity. Results of GO enrichment analysis showed that DEGs upregulated in C0 were significantly enriched in DNA replication pathway and cell cycle process as well (Figures 3C and 3D). Besides, several histone genes, such as *HIST1H4C*, *HIST1H1C*, exhibited higher expression levels in subpopulations C1 (Figure 3B). Contrarily, cells in C5 displayed aging characteristics, although proportion of which is very small in the total populations, and almost all these cells assigned to G1 phase belonging to UC1 sample (Figure S3B). We thought that cells in subpopulation C5 may have experienced mutation or replicative senescence during expansion and they were removed from the following analysis. Across those candidate subpopulations, several markers of MSCs showed similar expression level (Figure 3E). Meanwhile, we noted that collagen and chemokine genes across these subpopulations were differentially expressed. Specially, expression of collagen genes was much higher in C3 while expression of chemokine genes was higher in C4 (Figures S3C and S3D). Furthermore, we identified several candidate surface markers, which could be used to sort those subpopulations for further physiological and functional studies (Figure 3F).

**Figure 3.**
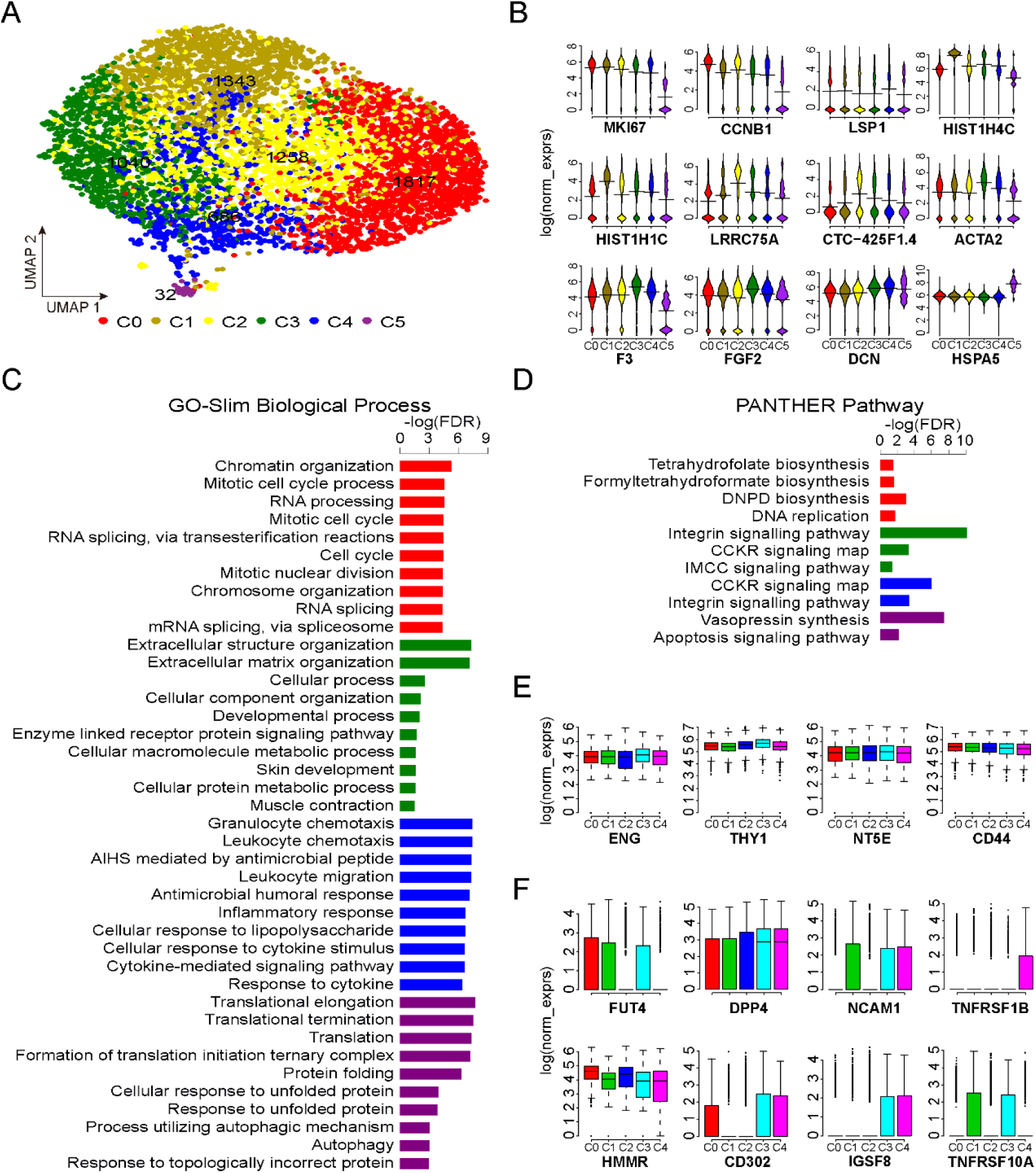
Candidate subpopulations with different functional characteristics. (A) UMAP visualizing the results of cell clustering. (B) Bean plots showing expression of several DEGs among the six subpopulations. (C) Pathways significantly enriched for the genes differentially expressed in one subpopulation compared to others. (D) GO-slim biological process enriched for the genes differentially expressed in one subpopulation compared to others (E) Boxplots showing expression of classic MSC marker genes in subpopulations. (F) Example of candidate markers showing different expression pattern among the five subpopulations (C0-C4). C0, red; C1, olive; C2, yellow; C3, green; C4, blue; C5, purple. IMCC: Inflammation mediated by chemokine and cytokine; DNPD: De novo pyrimidine deoxyribonucleotide.

In terms of MSC function, on which the MSC clinical application were theoretically based, the DEGs upregulated in subpopulations C3 were enriched in extracellular structure organization, developmental process, and muscle contraction, while DEGs upregulated in subpopulations C4 were associated with immunomodulation function (Figure 4A). Secretome analysis revealed that increased levels of some cytokines, such as CCL2, GCSF, VEGF, and IL-7, are positively correlated with immunosuppression (Chinnadurai et al., 2018). Among these subpopulation, expression levels of *CCL2* and *CSF3* are highest in C4 subpopulation (Figure S6F), implicating its immunomodulation therapeutic potential. Besides, lineage differentiation score among these subpopulations were different, indicating their distinct differentiation propensity to osteogenic, chondrogenic, adipogenic, myogenic or neurogenic cells (Figures 4B-4F).

**Fig 4.**
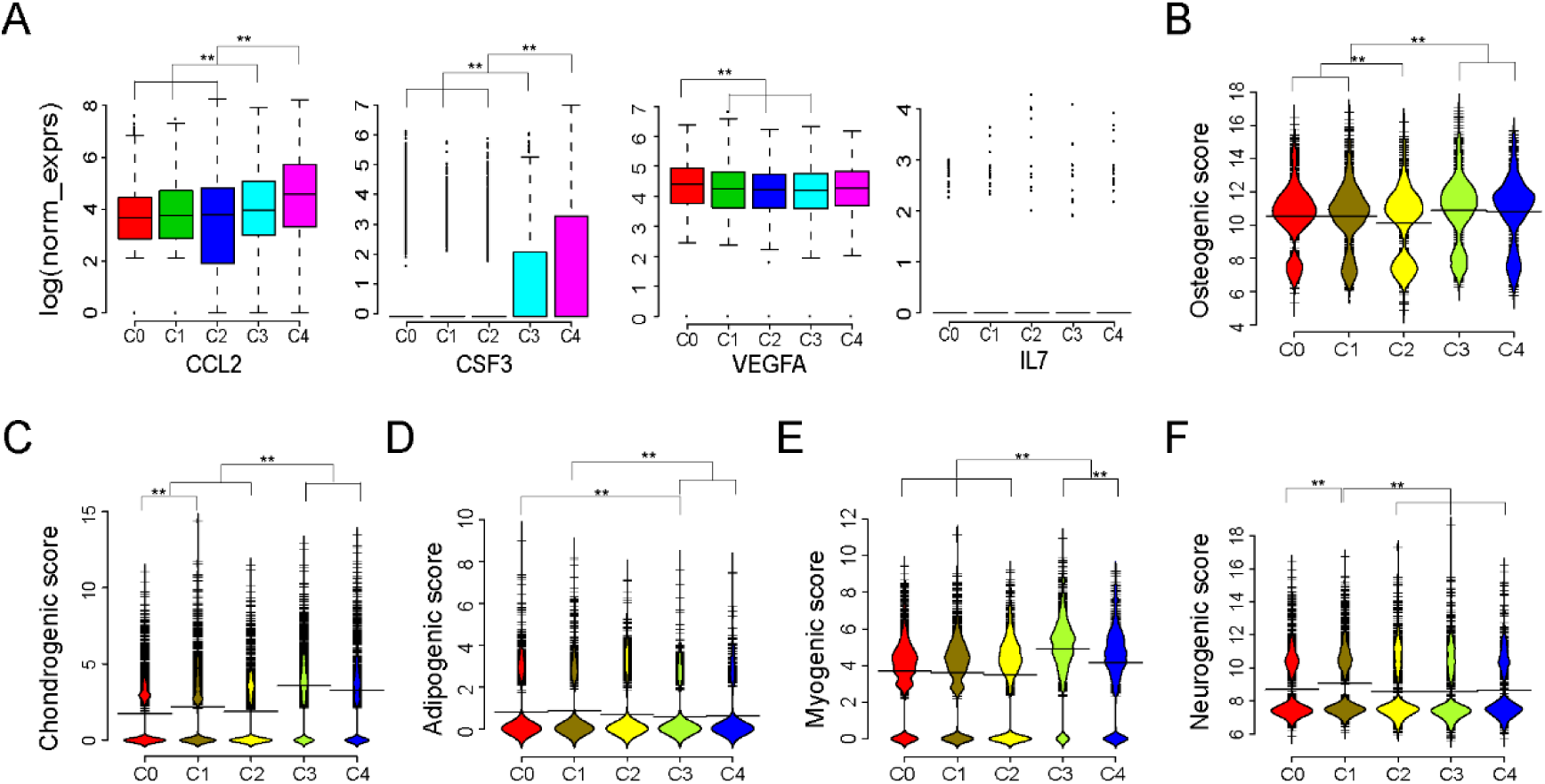
Candidate subpopulations showing different predicted potency on differentiation and immunosuppression. (A) Boxplots showing expression of genes correlated with PBMC suppression across the five candidate subpopulations (C0–C4). (B-F) Bean plots showing distribution of log (normalized expression) values of osteogenic score (B), chondrogenic score (C), adipogenic score (D), myogenic score (E), and neurogenic score (F) across the five candidate subpopulations (C0–C4). Wilcoxon Rank Sum test were performed for significant test, ** P < 0.001.

### Sing-cell Transcriptome Comparison between WJMSCs and ADMSCs

To provide insights into the heterogeneity of MSCs, several previous studies have compared gene expression of MSCs isolated from different sources using bulk-cell transcriptomic profiles (Alhattab et al., 2019; Fong et al., 2011; Lee et al., 2004; Ma et al., 2019; Meng et al., 2019; Taşkiran and Karaosmanoğlu, 2019). However, even MSCs derived from the same tissue exhibited inter-population functional heterogeneity, such as different differentiation potency and proliferation capacity. Bulk RNA-seq measures the average expression of genes, which is the sum of cell type-specific gene expression weighted by cell type proportions. Bulk transcriptome comparisons may hide some meaningful information that can help to elucidate the underlying mechanisms of functional heterogeneity. Thus, here we compared transcriptome data at the single-cell level between WJMSCs and ADMSCs. As expected, a lot of highly expressed classic MSCs surface markers are shared between these two MSCs, including *ENG, NT5E, THY1*, and *CD44* (Figures 5A, 5B and Table S1). Meanwhile, some unshared CDs were identified (Figure 5B), which suggest phenotypic diversity between WJMSCs and ADMSCs. These unshared genes involved in different cell signaling pathways inferred from pathway enrichment analysis of the top50 CDs (Figure 5C). Not surprisingly, some of these unshared CDs, although ranked in the top 50 genes by average expression (Figures 1A and 5A), only expressed at high levels in a small proportion of the MSCs (Figure 5D). Some of the unshared CDs are expressed (or not expressed) in majority of the MSCs derived from one tissue, but expressed only in a small proportion of the MSCs in the other one, such as *CD36*, which plays an important role in the formation of intracellular lipid droplets (Durandt et al., 2016), as well as *ITGA1*, *ITGA2*, and *PI16* (Figure 5D and Table S2). Those CD genes hold the potential to be used as markers for subpopulations sorting for further physiological and functional research. Accordingly, we also found that several markers, which have been reported to identify special MSC subpopulations with different biological functions, are expressed weakly in a small portion of MSCs, including *CD271* (*NGFR*) (Battula et al., 2009; Kohli et al., 2019), *CD146* (*MCAM*) (Espagnolle et al., 2014), *CXCR4*, *NES* (Morrison and Scadden, 2014), *CD106* (*VCAM1*) (Yang et al., 2013), except *PDGFRA*, which are highly expressed in most cells in WJMSC and ADMSC (Figure 5E and Table S2).

**Figure 5.**
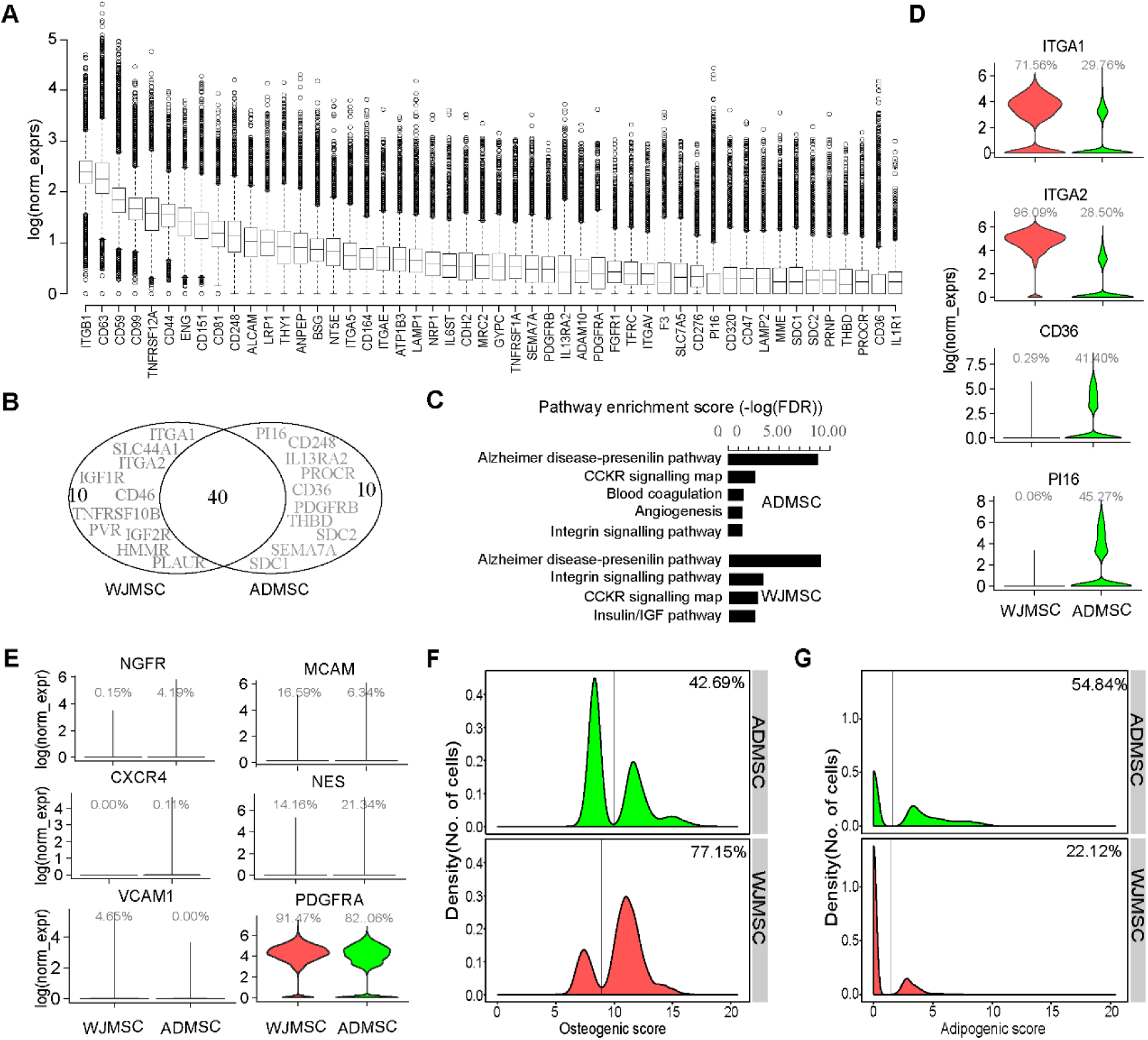
Transcriptome Comparison at the single-cell level between WJMSC and ADMSC. (A) Boxplot showing the top 50 CD genes ranked by average normalized expression in ADMSCs. (B) Venn diagram showing top 50 ADMSCs CD genes overlap with the top 50 WJMSC CD genes, unshared genes were highlighted. (C) Pathway enrichment of top 50 CD genes expressed in ADMSC and WJMSC. (D) Example of CD genes showing different expression percentage between ADMSC and WJMSC. (E) Violin plots showing reported MSC subpopulations in ADMSC and WJMSC. (F and G) Density distribution showing osteogenic score (F) and adipogenic score (G) between ADMSC and WJMSC. Percentage indicating proportion of cells assigned to the right side of the line.

Increasing reports demonstrated that MSCs derived from different sources exhibited distinct biological properties, such as proliferative capacity, multi-lineage differentiation ability, and immunomodulation potency (Via et al., 2012), although they all meet the minimal criteria for defining multipotent MSCs. As regards the differentiation ability, we evaluated it by calculating lineage differentiation score using single cell gene expression data from WJMSCs and ADMSCs. Interestingly, density distributions of lineage score displayed two major peaks, while only one peak was observed in housekeeping genes analysis (Figures. 5F, 5G and S4A-S4D), which indicate the existence of multiple subpopulations in MSCs. Density distribution of osteogenic score showed that 77.15 % of cells in WJMSCs had high osteogenic score vs. 42.69 % in ADMSCs (Figure. 5F). On the contrary, more cells (54.84 %) in ADMSCs have higher adipogenic score than in WJMSCs (22.12 %) (Figure 5G). These results suggested that WJMSCs have the propensity towards the osteogenic lineage while ADMSCs are inclined to differentiate into adipose cells, which are in line with previous studies (Li et al., 2014). Moreover, difference in other lineage differentiation potency are also existed, such as the chondrogenic and myogenic potential (Figures S4A and S4C).

Recently, human skeletal stem cell (SSC) and adipose progenitor cells were identified (Chan et al., 2018; Merrick and Sakers, 2019). Human SSC with PDPN ^+^/CD146 (MCAM) ^−^/CD73(NT5E) ^+^/CD164 ^+^ phenotype have the ability to generate progenitors of bone, cartilage, and stroma, but not fat (Chan et al., 2018). Human adipose progenitor cells expressed DPP4 are able to give rise to committed ICAM1^+^ preadipocytes (Merrick and Sakers, 2019). To determine whether cultured MSCs consist of cells with phenotype as SSC or adipose progenitors, we analyzed single cell expression data with above marker genes. Notably, proportion of cells expressed SSC markers in WJMSC is much higher than in ADMSC (Figures S4E and S4F), while more cells with adipose progenitor’s markers exist in ADMSC (Figures S4G and S4H). These results further indicated that bulk-cell variations among MSCs from different sources may originate from composition diversity of distinct subpopulations.

## DISCUSSION

MSCs are promising cell therapy products with great potential in promoting tissue regeneration and modulating inflammation. However, significant variations were reported during culturing MSCs that were isolated from different donors and different tissue sites. Unrefined, non-standardized isolation and culture techniques become the challenges of standardization in processes of MSC products manufacturing and quality management. Even in a “homogeneous” population, which defined by the classic minimal criteria, including cell size (Majore et al., 2009), morphology (Klinker et al., 2017; Marklein et al., 2019; Marklein et al., 2016), proliferation capacity (Majore et al., 2009), differentiation potency (Russell et al., 2010), and immunomodulation capacity (Klinker et al., 2017; Marklein et al., 2019), these cells still display phenotype and function heterogeneity among individual cells (Pevsner-Fischer et al., 2011). In previous clinical trials, functional variation and heterogeneity in MSCs are potentially the main reasons lead to inconsistent or controversial results (Phinney, 2012; Zhang et al., 2015). The underlying molecular mechanisms that lead to MSCs functional variation and heterogeneity at the cell population level remain unknown, which require further investigation and elucidation.

Recently, several studies have been performed to investigated into the heterogeneity of cultured MSCs by single cell transcriptomic analysis (Huang et al., 2019; Khong et al., 2019; Liu et al., 2019; Wang et al., 2019). Huang et al. profiled the transcriptomes of 361 single MSCs derived from two umbilical cords (UC-MSCs) that were harvested at different passages and stimulated with or without inflammatory cytokines. Following analysis, they concluded that *in vitro* expanded UC-MSCs are a well-organized population with limited heterogeneity, which is mainly caused by distinct distribution in cell cycle phases (Huang et al., 2019). However, the number of cells sequenced for each condition is small (~50 cells per condition), and they did not remove the cell cycle effects for the subpopulations identification. Besides, the only one marker (HMMR) they used to sort the cells to confirm their hypothesis may be unable to isolate different subpopulations. Liu et al. performed a large-scale single-cell transcriptomic sequencing of 24 370 cultured ADMSCs from three donors (Liu et al., 2019). They regressed out batch and cell cycle effects before candidate subpopulation classification, however, the results they exhibited in the report were limited to the data analysis pipeline. Wang et al. sequenced a total of 103 single hWJMSCs from three umbilical cords and 63 single hBMMSCs cells from two different donors, and just focused on gene expression comparison between the two different sources derived MSCs (Wang et al., 2019). Thus, the cellular transcriptomic heterogeneity within a MSC population cultured *in vitro* still have not been comprehensively investigated at the single-cell level.

In this study, we dissected gene-expression heterogeneity of human primary WJMSCs cultured *in vitro* using scRNA-seq. Single-cell RNA sequencing technologies can offer an unbiased approach for understanding the extent, basis and function of gene expression variation between seemingly identical cells, revealing complex and rare cell populations, uncovering regulatory relationships between genes, and tracking the trajectories of distinct cell lineages in development (Gurtner et al., 2018; Tang et al., 2009). In primary WJMSCs, we found that the HVGs are significantly enriched in extracellular region with binding function, involved in developmental process, signal transduction, cell proliferation, etc. For example, MKI67, a marker of proliferation, were identified as one of the HVGs, implying different proliferate capacity among individual cells. In terms of therapeutic potential, these genes are associated with functional characteristics of MSCs, such as integrin signaling pathway, angiogenesis, and inflammation mediated by chemokine and cytokine signaling pathway (Figure 2E). Integrin signaling pathway plays a critical role in homing of MSCs to bone, osteogenic differentiation, and bone formation, and even some integrins are suggested as targets to promote bone formation and repair (Di Maggio et al., 2017; Marie, 2013; Olivares-Navarrete et al., 2015). Several integrin genes were identified in our data with highly variable expression across the cells, such as *ITGA5* and *ITGB1*, which respectively encode α5 and β1 and together form the α5β1 integrin, a cell-surface receptor for fibronectin implicated in the control of osteoblastogenesis (Marie, 2013; Park et al., 2011). Pro-angiogenesis is one of the important biological properties of MSCs, implicating in promoting wound healing and tissue repair (Watt et al., 2013; Wu et al., 2007). Genes related to the angiogenesis, such as *ANGPT1*, *PDGFRA*, *VEGFA*, etc., were identified in our HVGs (Table S3). Studies have reported that *ANGPT1* gene-modified human MSCs could promote angiogenesis and reduce acute pancreatitis in rats (Hua et al., 2014), while *PDGFRA*^+^ MSCs have enhanced skin repair/regeneration potential (Iinuma et al., 2015). *VEGFA*, and other two cytokines, *CXCL5* and *CXCL8* (IL-8), were required for the angiogenic activity of MSCs and have been selected as an assay matrix for angiogenic potency assay for MultiStem product (Galipeau et al., 2016; Lehman et al., 2012). Furthermore, *in vitro* co-culture assays demonstrated that the increased levels of *VEGFA* and chemokine *CCL2* in MSCs were positively correlated with PBMC suppression (Chinnadurai et al., 2018). Chemokines, a family of small cytokines, are recognized as key mediators of MSCs migration and immunosuppression (Hocking, 2015; Ren et al., 2008). Notably, most of the chemokines expressed in primary WJMSCs were highly heterogeneous, including above mentioned *CCL2*, *CXCL5*, and *CXCL8*. These results indicated that highly variable genes within WJMSCs are associated with classic MSC functional properties, and suggested the existence of potential subpopulations with different gene expression patterns.

Although cultured MSCs meet the minimal criteria with classic phenotype, increasing reports demonstrated that many cell surface membrane proteins are not uniformly expressed in MSC (Mo et al., 2016). Several subpopulations with different phenotype, property and therapeutic potential have been identified in MSCs derived from different tissues. Some subpopulations express *CXCR4* and have a propensity to migrate to sites of tissue injury (Cheng et al., 2008), while some express *VCAM-1* (*CD106*) and show priority in immunosuppression (Yang et al., 2013). However, these above reported markers only weakly expressed in WJMSCs (Figure 4E), may unable to serve as effective markers to isolated these subpopulations in WJMSCs. Here, we classified WJMSCs into several candidate subpopulations (C0-C5) with different functional characteristics. Among these candidate subpopulations, DEG analysis indicated that C0 and C1 show greater proliferation ability while C3 and C4 have greater osteogenic and chondrogenic differentiation potency. Myogenic score is also significantly higher in C3, implying its potential in myocardial repair application. As to immunomodulation, we found most of the chemokines and some immune-related cytokines detected in WJMSCs upregulate in C4. Taken together, these candidate subpopulations identified in primary WJMSC would be valuable for further biological characterization via experimental investigations and clinical researches.

More and more clinical trials are being using MSCs to treat diverse diseases these days. However, challenges of developing potency assays for MSC-like products hinder their clinical applications, which include variability of tissue sources, largely undefined mechanisms of action in humans, and lack of reference standards (Chinnadurai et al., 2018; Galipeau and Krampera, 2015; Galipeau et al., 2016; Hematti, 2016). By comparing gene expression between WJMSCs and ADMSCs at the single-cell level, we found that HVGs identified in WJMSC are largely overlapped with ADMSC (Figure S5A), though there are unshared HVGs that might contribute to the differences in lineage potential. Furthermore, functional enrichment analysis of HVGs from ADMSC showed similar results as those from WJMSC (Figures S5B-S5G), suggesting that these HVGs play critical roles in MSCs, and may serve as candidate markers for further potency association studies.

In summary, highly variable genes within MSCs are significantly enriched in extracellular region with binding function, involved in developmental process, signal transduction, cell proliferation. Regarding therapeutic potential, these genes are associated with many functional characteristics of MSCs, including inflammation mediated by chemokine and cytokine signaling, integrin signaling, and angiogenesis. Candidate subpopulations identified in MSCs also show different functional characteristics, such as proliferation, differentiation propensity, and immunosuppression potency. Further studies in cell-to-cell variability in transcriptome, proteome, secretome, and epigenome on MSCs derived from different tissues, will increase our understanding of the heterogeneity associated with MSC function and facilitate the development of MSC release criteria for clinical application.

## EXPERIMENTAL PROCEDURES

### Cell Isolation and Culture

This study was approved by the Ethics Committee of BGI-IRB. Human umbilical cord tissue were collected from naturally delivered full-term newborns (n = 3, two females and one male). WJs were isolated from umbilical cord after dissection and mechanically dissociated into tissue explants of approximately 2 mm^2^, which were then seeded into T75 flasks and cultured in UltraCULTURETM Serum-free Medium (LONZA) at 37 °C with 5 % CO_2_ in a humidified atmosphere.. After cell density reached about 80 % confluence, cells were dissociated with TrypLE™ Select (ThermoFisher Scientific) incubated at 37 °C for 5 min. The collected cells were immediately used for single cell library construction, sub-cultured into a new culture dish, tri-lineage differentiation potency test, or freezing in liquid nitrogen for long-term banking.

### scRNA-seq and Analysis

scRNA-seq experiment was performed using the Chromium Single Cell Gene Expression Solution, V2 Chemistry (10x Genomics), following the manufacturer’s protocol. Briefly, the collected cells were washed with PBS twice, and resuspended in 500 μl PBS, targeting the required 500 cells/μl concentration. We pipetted 6.4 μl cell suspension, targeting the recovery of about 2 000 cells per sample. Single-cell RNA-seq libraries were obtained following the 10x Genomics recommended protocol, using the reagents included in the Chromium Single Cell 3′ v2 Reagent Kit. Libraries were sequenced on the BGISEQ-500 (BGI) instrument (Natarajan et al., 2019) using 26 cycles (cell barcode and UMI (Islam et al., 2014)) for read1 and 108 cycles (sample index and transcript 3’ end) for read2, obtaining about 5 × 10^8^ raw paired reads.

scRNA-seq data analysis was available in the Supplemental Experimental Procedures for details. Briefly, the scRNA-seq data was processed using cellranger-2.0.0 for mapping. Outlier cells using the median absolute deviation from the median total library size (logarithmic scale), total gene numbers (logarithmic scale), as well as mitochondrial percentage, as implemented in scran package, using a cutoff of 3 (Lun et al., 2016). Any gene expressed across all the cells by average UMI less than 0.1 was removed. Cell cycle phase assignment and removal, highly variable genes identification, linear and nonlinear dimension reduction, clustering and differential expression analysis were performed using Seurat package (Butler et al., 2018).

### Functional Enrichment Analysis

GO-slim, protein class and pathways overrepresentation Test enrichment analyses were performed using PANTHER™ Version 14.1 according to Mi et al. (Mi et al., 2019) via test type of Fisher’s Exact, applying the Benjamini–Hochberg false discovery rate (FDR) correction for multiple testing.

### Lineage Differentiation Potency Evaluation

Using the marker genes listed in Supplementary Table S5, we calculated osteogenic, adipogenic, chondrogenic, neurogenic, and myogenic ‘scores’ according to (Schwalie et al., 2018). Specifically, Score defined as a single numeric value representative of the expression of multiple marker genes, the sum of log normalized expression across all markers in a category. Housekeeping genes were also used and named as housekeeping score.

## ACCESSION NUMBERS

The data that support the findings of this study have been deposited in the CNSA (https://db.cngb.org/cnsa/) of CNGBdb with accession number CNP0000562.

## SUPPLEMENTAL INFORMATION

Supplemental Information includes Supplemental Experimental Procedures, five Figures, and five tables.

## AUTHOR CONTRIBUTIONS

Conceptualization, C.S., and X.Z.; Methodology and Investigation, C.S., L.W., and T.H. Writing, C.S. and H.W.; Funding Acquisition, X.Z.

## ACKNOWLEDGMENTS

This work was supported by Shenzhen Municipal Government of China (No. CYFWT201507021005 to X.Z.) and Science, Technology and Innovation Commission of Shenzhen Municipality under grant No. KQJSCX20170322143848 413 to X.Z. The funders had no role in study design, data collection and analysis, decision to publish, or preparation of the manuscript. It was also supported by BGI-Shenzhen. The funder provided support in the form of salaries for C.S., L.W., T.H., and X.Z., but did not have any additional role in the study design, data collection and analysis, decision to publish, or preparation of the manuscript. We thank members of China National GeneBank for technical support.

## REFERENCES

Abdallah, B.M., and Kassem, M. (2008). Human mesenchymal stem cells: from basic biology to clinical applications. Gene Ther 15, 109–116.

Aggarwal, S., and Pittenger, M.F. (2005). Human mesenchymal stem cells modulate allogeneic immune cell responses. Blood 105, 1815–1822.

Alhattab, D., Jamali, F., Ali, D., Hammad, H., Adwan, S., Rahmeh, R., Samarah, O., Salah, B., Hamdan, M., and Awidi, A. (2019). An insight into the whole transcriptome profile of four tissue-specific human mesenchymal stem cells. Regen Med.

Baksh, D., Song, L., and Tuan, R.S. (2004). Adult mesenchymal stem cells: characterization, differentiation, and application in cell and gene therapy. J Cell Mol Med 8, 301–316.

Battula, V.L., Treml, S., Bareiss, P.M., Gieseke, F., Roelofs, H., de Zwart, P., Muller, I., Schewe, B., Skutella, T., Fibbe, W.E., et al. (2009). Isolation of functionally distinct mesenchymal stem cell subsets using antibodies against CD56, CD271, and mesenchymal stem cell antigen-1. Haematologica 94, 173–184.

Bustos, M.L., Huleihel, L., Kapetanaki, M.G., Lino-Cardenas, C.L., Mroz, L., Ellis, B.M., McVerry, B.J., Richards, T.J., Kaminski, N., Cerdenes, N., et al. (2014). Aging mesenchymal stem cells fail to protect because of impaired migration and antiinflammatory response. Am J Respir Crit Care Med 189, 787–798.

Butler, A., Hoffman, P., Smibert, P., Papalexi, E., and Satija, R. (2018). Integrating single-cell transcriptomic data across different conditions, technologies, and species. Nat Biotechnol 36, 411–420.

Chan, C.K.F., Gulati, G.S., Sinha, R., Tompkins, J.V., Lopez, M., Carter, A.C., Ransom, R.C., Reinisch, A., Wearda, T., Murphy, M., et al. (2018). Identification of the Human Skeletal Stem Cell. Cell 175, 43–56.e21.

Chen, S.L., Fang, W.W., Ye, F., Liu, Y.H., Qian, J., Shan, S.J., Zhang, J.J., Chunhua, R.Z., Liao, L.M., Lin, S., et al. (2004). Effect on left ventricular function of intracoronary transplantation of autologous bone marrow mesenchymal stem cell in patients with acute myocardial infarction. Am J Cardiol 94, 92–95.

Cheng, Z., Ou, L., Zhou, X., Li, F., Jia, X., Zhang, Y., Liu, X., Li, Y., Ward, C.A., Melo, L.G., et al. (2008). Targeted migration of mesenchymal stem cells modified with CXCR4 gene to infarcted myocardium improves cardiac performance. Mol Ther 16, 571–579.

Chinnadurai, R., Rajan, D., Qayed, M., Arafat, D., Garcia, M., Liu, Y., Kugathasan, S., Anderson, L.J., Gibson, G., and Galipeau, J. (2018). Potency Analysis of Mesenchymal Stromal Cells Using a Combinatorial Assay Matrix Approach. Cell reports 22, 2504–2517.

Di Maggio, N., Martella, E., Frismantiene, A., Resink, T.J., Schreiner, S., Lucarelli, E., Jaquiery, C., Schaefer, D.J., Martin, I., and Scherberich, A. (2017). Extracellular matrix and alpha5beta1 integrin signaling control the maintenance of bone formation capacity by human adipose-derived stromal cells. Sci Rep 7, 44398.

Docheva, D., Popov, C., Mutschler, W., and Schieker, M. (2007). Human mesenchymal stem cells in contact with their environment: surface characteristics and the integrin system. J Cell Mol Med 11, 21–38.

Dominici, M., Le Blanc, K., Mueller, I., Slaper-Cortenbach, I., Marini, F., Krause, D., Deans, R., Keating, A., Prockop, D., and Horwitz, E. (2006). Minimal criteria for defining multipotent mesenchymal stromal cells. The International Society for Cellular Therapy position statement. Cytotherapy 8, 315–317.

Durandt, C., van Vollenstee, F.A., Dessels, C., Kallmeyer, K., de Villiers, D., Murdoch, C., Potgieter, M., and Pepper, M.S. (2016). Novel flow cytometric approach for the detection of adipocyte subpopulations during adipogenesis. J Lipid Res 57, 729–742.

Espagnolle, N., Guilloton, F., Deschaseaux, F., Gadelorge, M., Sensebe, L., and Bourin, P. (2014). CD146 expression on mesenchymal stem cells is associated with their vascular smooth muscle commitment. J Cell Mol Med 18, 104–114.

Even-Ram, S., Artym, V., and Yamada, K.M. (2006). Matrix control of stem cell fate. Cell 126, 645–647.

Fong, C.Y., Chak, L.L., Biswas, A., Tan, J.H., Gauthaman, K., Chan, W.K., and Bongso, A. (2011). Human Wharton’s jelly stem cells have unique transcriptome profiles compared to human embryonic stem cells and other mesenchymal stem cells. Stem cell reviews 7, 1–16.

Frantz, C., Stewart, K.M., and Weaver, V.M. (2010). The extracellular matrix at a glance. J Cell Sci 123, 4195–4200.

Fukuchi, Y., Nakajima, H., Sugiyama, D., Hirose, I., Kitamura, T., and Tsuji, K. (2004). Human placenta-derived cells have mesenchymal stem/progenitor cell potential. Stem Cells 22, 649–658.

Galipeau, J., and Krampera, M. (2015). The challenge of defining mesenchymal stromal cell potency assays and their potential use as release criteria. Cytotherapy 17, 125–127.

Galipeau, J., Krampera, M., Barrett, J., Dazzi, F., Deans, R.J., DeBruijn, J., Dominici, M., Fibbe, W.E., Gee, A.P., Gimble, J.M., et al. (2016). International Society for Cellular Therapy perspective on immune functional assays for mesenchymal stromal cells as potency release criterion for advanced phase clinical trials. Cytotherapy 18, 151–159.

Galipeau, J., and Sensebe, L. (2018). Mesenchymal Stromal Cells: Clinical Challenges and Therapeutic Opportunities. Cell stem cell 22, 824–833.

Ghannam, S., Bouffi, C., Djouad, F., Jorgensen, C., and Noel, D. (2010). Immunosuppression by mesenchymal stem cells: mechanisms and clinical applications. Stem Cell Res Ther 1, 2.

Guilak, F., Cohen, D.M., Estes, B.T., Gimble, J.M., Liedtke, W., and Chen, C.S. (2009). Control of stem cell fate by physical interactions with the extracellular matrix. Cell stem cell 5, 17–26.

Gurtner, G.C., Hwang, B., Lee, J.H., and Bang, D. (2018). Single-cell RNA sequencing technologies and bioinformatics pipelines. Stem Cells 50, 96.

Hafemeister, C., and Satija, R. (2019). Normalization and variance stabilization of single-cell RNA-seq data using regularized negative binomial regression. bioRxiv, 576827.

Hamidouche, Z., Fromigue, O., Ringe, J., Haupl, T., Vaudin, P., Pages, J.C., Srouji, S., Livne, E., and Marie, P.J. (2009). Priming integrin alpha5 promotes human mesenchymal stromal cell osteoblast differentiation and osteogenesis. Proc Natl Acad Sci U S A 106, 18587–18591.

Hematti, P. (2016). Characterization of mesenchymal stromal cells: potency assay development. Transfusion (Paris) 56, 32s–35s.

Hocking, A.M. (2015). The Role of Chemokines in Mesenchymal Stem Cell Homing to Wounds. Adv Wound Care (New Rochelle) 4, 623–630.

Hua, J., He, Z.G., Qian, D.H., Lin, S.P., Gong, J., Meng, H.B., Yang, T.S., Sun, W., Xu, B., Zhou, B., et al. (2014). Angiopoietin-1 gene-modified human mesenchymal stem cells promote angiogenesis and reduce acute pancreatitis in rats. Int J Clin Exp Pathol 7, 3580–3595.

Huang, Y., Li, Q., Zhang, K., Hu, M., Wang, Y., Du, L., and Lin, L. (2019). Single cell transcriptomic analysis of human mesenchymal stem cells reveals limited heterogeneity. 10, 368.

Iinuma, S., Aikawa, E., Tamai, K., Fujita, R., Kikuchi, Y., Chino, T., Kikuta, J., McGrath, J.A., Uitto, J., Ishii, M., et al. (2015). Transplanted bone marrow-derived circulating PDGFRalpha+ cells restore type VII collagen in recessive dystrophic epidermolysis bullosa mouse skin graft. J Immunol 194, 1996–2003.

Ikebe, C., and Suzuki, K. (2014). Mesenchymal stem cells for regenerative therapy: optimization of cell preparation protocols. BioMed research international 2014, 951512.

Islam, S., Zeisel, A., Joost, S., La Manno, G., Zajac, P., Kasper, M., Lonnerberg, P., and Linnarsson, S. (2014). Quantitative single-cell RNA-seq with unique molecular identifiers. Nature methods 11, 163–166.

Jin, H.J., Bae, Y.K., Kim, M., Kwon, S.J., Jeon, H.B., Choi, S.J., Kim, S.W., Yang, Y.S., Oh, W., and Chang, J.W. (2013). Comparative analysis of human mesenchymal stem cells from bone marrow, adipose tissue, and umbilical cord blood as sources of cell therapy. International journal of molecular sciences 14, 17986–18001.

Karussis, D., Karageorgiou, C., Vaknin-Dembinsky, A., Gowda-Kurkalli, B., Gomori, J.M., Kassis, I., Bulte, J.W., Petrou, P., Ben-Hur, T., Abramsky, O., et al. (2010). Safety and immunological effects of mesenchymal stem cell transplantation in patients with multiple sclerosis and amyotrophic lateral sclerosis. Arch Neurol 67, 1187–1194.

Kharaziha, P., Hellstrom, P.M., Noorinayer, B., Farzaneh, F., Aghajani, K., Jafari, F., Telkabadi, M., Atashi, A., Honardoost, M., Zali, M.R., et al. (2009). Improvement of liver function in liver cirrhosis patients after autologous mesenchymal stem cell injection: a phase I-II clinical trial. Eur J Gastroenterol Hepatol 21, 1199–1205.

Khong, S.M.L., Lee, M., Kosaric, N., Khong, D.M., Dong, Y., Hopfner, U., Aitzetmuller, M.M., Duscher, D., and Schafer, R. (2019). Single-Cell Transcriptomics of Human Mesenchymal Stem Cells Reveal Age-Related Cellular Subpopulation Depletion and Impaired Regenerative Function. 37, 240–246.

Klinker, M.W., Marklein, R.A., Lo Surdo, J.L., Wei, C.H., and Bauer, S.R. (2017). Morphological features of IFN-gamma-stimulated mesenchymal stromal cells predict overall immunosuppressive capacity. 114, E2598–e2607.

Kohli, N., Al-Delfi, I.R.T., Snow, M., Sakamoto, T., Miyazaki, T., Nakajima, H., Uchida, K., and Johnson, W.E.B. (2019). CD271-selected mesenchymal stem cells from adipose tissue enhance cartilage repair and are less angiogenic than plastic adherent mesenchymal stem cells. Sci Rep 9, 3194.

Krampera, M., Pizzolo, G., Aprili, G., and Franchini, M. (2006). Mesenchymal stem cells for bone, cartilage, tendon and skeletal muscle repair. Bone 39, 678–683.

Le Blanc, K., Frassoni, F., Ball, L., Locatelli, F., Roelofs, H., Lewis, I., Lanino, E., Sundberg, B., Bernardo, M.E., Remberger, M., et al. (2008). Mesenchymal stem cells for treatment of steroid-resistant, severe, acute graft-versus-host disease: a phase II study. Lancet 371, 1579–1586.

Lee, R.H., Kim, B., Choi, I., Kim, H., Choi, H.S., Suh, K., Bae, Y.C., and Jung, J.S. (2004). Characterization and expression analysis of mesenchymal stem cells from human bone marrow and adipose tissue. Cell Physiol Biochem 14, 311–324.

Lehman, N., Cutrone, R., Raber, A., Perry, R., Van’t Hof, W., Deans, R., Ting, A.E., and Woda, J. (2012). Development of a surrogate angiogenic potency assay for clinical-grade stem cell production. Cytotherapy 14, 994–1004.

Li, X., Bai, J., Ji, X., Li, R., Xuan, Y., and Wang, Y. (2014). Comprehensive characterization of four different populations of human mesenchymal stem cells as regards their immune properties, proliferation and differentiation. Int J Mol Med 34, 695–704.

Liu, X., Xiang, Q., Xu, F., Huang, J., Yu, N., Zhang, Q., Long, X., and Zhou, Z. (2019). Single-cell RNA-seq of cultured human adipose-derived mesenchymal stem cells. Scientific data 6, 190031.

Lun, A.T., McCarthy, D.J., and Marioni, J.C. (2016). A step-by-step workflow for low-level analysis of single-cell RNA-seq data with Bioconductor. F1000Research 5, 2122.

Ma, J., Wu, J., Han, L., Jiang, X., Yan, L., Hao, J., and Wang, H. (2019). Comparative analysis of mesenchymal stem cells derived from amniotic membrane, umbilical cord, and chorionic plate under serum-free condition. Stem Cell Res Ther 10, 19.

Macosko, E.Z., Basu, A., Satija, R., Nemesh, J., Shekhar, K., Goldman, M., Tirosh, I., Bialas, A.R., Kamitaki, N., Martersteck, E.M., et al. (2015). Highly Parallel Genome-wide Expression Profiling of Individual Cells Using Nanoliter Droplets. Cell 161, 1202–1214.

Majore, I., Moretti, P., Hass, R., and Kasper, C. (2009). Identification of subpopulations in mesenchymal stem cell-like cultures from human umbilical cord. Science 7, 6.

Marie, P.J. (2013). Targeting integrins to promote bone formation and repair. Nat Rev Endocrinol 9, 288–295.

Marklein, R.A., Klinker, M.W., Drake, K.A., Polikowsky, H.G., Lessey-Morillon, E.C., and Bauer, S.R. (2019). Morphological profiling using machine learning reveals emergent subpopulations of interferon-gamma-stimulated mesenchymal stromal cells that predict immunosuppression. Cytotherapy 21, 17–31.

Marklein, R.A., Lo Surdo, J.L., Bellayr, I.H., Godil, S.A., Puri, R.K., and Bauer, S.R. (2016). High Content Imaging of Early Morphological Signatures Predicts Long Term Mineralization Capacity of Human Mesenchymal Stem Cells upon Osteogenic Induction. Stem Cells 34, 935–947.

Mazzini, L., Ferrero, I., Luparello, V., Rustichelli, D., Gunetti, M., Mareschi, K., Testa, L., Stecco, A., Tarletti, R., Miglioretti, M., et al. (2010). Mesenchymal stem cell transplantation in amyotrophic lateral sclerosis: A Phase I clinical trial. Proc Natl Acad Sci U S A 223, 229–237.

Meng, X., Sun, B., and Xiao, Z. (2019). Comparison in transcriptome and cytokine profiles of mesenchymal stem cells from human umbilical cord and cord blood. Gene 696, 10–20.

Merrick, D., and Sakers, A. (2019). Identification of a mesenchymal progenitor cell hierarchy in adipose tissue. 364.

Mi, H., Muruganujan, A., Huang, X., Ebert, D., Mills, C., Guo, X., and Thomas, P.D. (2019). Protocol Update for large-scale genome and gene function analysis with the PANTHER classification system (v.14.0). Nat Protoc 14, 703–721.

Mo, M., Wang, S., Zhou, Y., Li, H., and Wu, Y. (2016). Mesenchymal stem cell subpopulations: phenotype, property and therapeutic potential. Cell Mol Life Sci 73, 3311–3321.

Morrison, S.J., and Scadden, D.T. (2014). The bone marrow niche for haematopoietic stem cells. Nature 505, 327–334.

Natarajan, K.N., Miao, Z., Jiang, M., Huang, X., Zhou, H., Xie, J., Wang, C., Qin, S., Zhao, Z., Wu, L., et al. (2019). Comparative analysis of sequencing technologies for single-cell transcriptomics. Genome Biol 20, 70.

Olivares-Navarrete, R., Hyzy, S.L., Park, J.H., Dunn, G.R., Haithcock, D.A., Wasilewski, C.E., Boyan, B.D., and Schwartz, Z. (2011). Mediation of osteogenic differentiation of human mesenchymal stem cells on titanium surfaces by a Wnt-integrin feedback loop. Biomaterials 32, 6399–6411.

Olivares-Navarrete, R., Rodil, S.E., Hyzy, S.L., Dunn, G.R., Almaguer-Flores, A., Schwartz, Z., and Boyan, B.D. (2015). Role of integrin subunits in mesenchymal stem cell differentiation and osteoblast maturation on graphitic carbon-coated microstructured surfaces. Biomaterials 51, 69–79.

Parekkadan, B., and Milwid, J.M. (2010). Mesenchymal stem cells as therapeutics. Annual review of biomedical engineering 12, 87–117.

Park, S.J., Gadi, J., Cho, K.W., Kim, K.J., Kim, S.H., Jung, H.S., and Lim, S.K. (2011). The forkhead transcription factor Foxc2 promotes osteoblastogenesis via up-regulation of integrin beta1 expression. Bone 49, 428–438.

Pevsner-Fischer, M., Levin, S., and Zipori, D. (2011). The origins of mesenchymal stromal cell heterogeneity. Stem cell reviews 7, 560–568.

Phinney, D.G. (2012). Functional heterogeneity of mesenchymal stem cells: implications for cell therapy. J Cell Biochem 113, 2806–2812.

Pijuan Sala, B., Diamanti, E., Shepherd, M., Laurenti, E., Wilson, N.K., Kent, D.G., Gottgens, B., Stuart, T., Butler, A., Hoffman, P., et al. (2019). Comprehensive Integration of Single-Cell Data. Blood 177, 1888–1902.e1821.

Prockop, D.J. (1997). Marrow stromal cells as stem cells for nonhematopoietic tissues. Science 276, 71–74.

Ranganath, S.H., Levy, O., Inamdar, M.S., and Karp, J.M. (2012). Harnessing the mesenchymal stem cell secretome for the treatment of cardiovascular disease. Cell stem cell 10, 244–258.

Ren, G., Zhang, L., Zhao, X., Xu, G., Zhang, Y., Roberts, A.I., Zhao, R.C., and Shi, Y. (2008). Mesenchymal stem cell-mediated immunosuppression occurs via concerted action of chemokines and nitric oxide. Cell stem cell 2, 141–150.

Romanov, Y.A., Svintsitskaya, V.A., and Smirnov, V.N. (2003). Searching for alternative sources of postnatal human mesenchymal stem cells: candidate MSC-like cells from umbilical cord. Stem Cells 21, 105–110.

Rozario, T., and DeSimone, D.W. (2010). The extracellular matrix in development and morphogenesis: a dynamic view. Dev Biol 341, 126–140.

Russell, K.C., Phinney, D.G., Lacey, M.R., Barrilleaux, B.L., Meyertholen, K.E., and O’Connor, K.C. (2010). In vitro high-capacity assay to quantify the clonal heterogeneity in trilineage potential of mesenchymal stem cells reveals a complex hierarchy of lineage commitment. Stem Cells 28, 788–798.

Samsonraj, R.M., Rai, B., Sathiyanathan, P., Puan, K.J., Rotzschke, O., Hui, J.H., Raghunath, M., Stanton, L.W., Nurcombe, V., and Cool, S.M. (2015). Establishing criteria for human mesenchymal stem cell potency. Stem Cells 33, 1878–1891.

Schwalie, P.C., Dong, H., Zachara, M., Russeil, J., Alpern, D., Akchiche, N., Caprara, C., Sun, W., Schlaudraff, K.U., Soldati, G., et al. (2018). A stromal cell population that inhibits adipogenesis in mammalian fat depots. Nature 559, 103–108.

Sethe, S., Scutt, A., and Stolzing, A. (2006). Aging of mesenchymal stem cells. Ageing research reviews 5, 91–116.

Tang, F., Barbacioru, C., Wang, Y., Nordman, E., Lee, C., Xu, N., Wang, X., Bodeau, J., Tuch, B.B., Siddiqui, A., et al. (2009). mRNA-Seq whole-transcriptome analysis of a single cell. Nature methods 6, 377–382.

Taşkiran, E.Z., and Karaosmanoğlu, B. (2019). Transcriptome analysis reveals differentially expressed genes between human primary bone marrow mesenchymal stem cells and human primary dermal fibroblasts. 43, 21–27.

Troyer, D.L., and Weiss, M.L. (2008). Wharton’s jelly-derived cells are a primitive stromal cell population. Stem Cells 26, 591–599.

Via, A.G., Frizziero, A., and Oliva, F. (2012). Biological properties of mesenchymal Stem Cells from different sources. Muscles, ligaments and tendons journal 2, 154–162.

Wakitani, S., Imoto, K., Yamamoto, T., Saito, M., Murata, N., and Yoneda, M. (2002). Human autologous culture expanded bone marrow mesenchymal cell transplantation for repair of cartilage defects in osteoarthritic knees. Osteoarthritis Cartilage 10, 199–206.

Wang, H.S., Hung, S.C., Peng, S.T., Huang, C.C., Wei, H.M., Guo, Y.J., Fu, Y.S., Lai, M.C., and Chen, C.C. (2004). Mesenchymal stem cells in the Wharton’s jelly of the human umbilical cord. Stem Cells 22, 1330–1337.

Wang, Y., Barrett, A.N., Fong, C.Y., Subramanian, A., Liu, W., Feng, Y., Choolani, M., Biswas, A., Rajapakse, J.C., and Bongso, A. (2019). Human Wharton’s Jelly Mesenchymal Stem Cells Show Unique Gene Expression Compared with Bone Marrow Mesenchymal Stem Cells Using Single-Cell RNA-Sequencing. Cell Death Dis 28, 196–211.

Watt, S.M., Gullo, F., van der Garde, M., Markeson, D., Camicia, R., Khoo, C.P., and Zwaginga, J.J. (2013). The angiogenic properties of mesenchymal stem/stromal cells and their therapeutic potential. Br Med Bull 108, 25–53.

Wu, Y., Chen, L., Scott, P.G., and Tredget, E.E. (2007). Mesenchymal stem cells enhance wound healing through differentiation and angiogenesis. Stem Cells 25, 2648–2659.

Yang, Z.X., Han, Z.B., Ji, Y.R., Wang, Y.W., Liang, L., Chi, Y., Yang, S.G., Li, L.N., Luo, W.F., Li, J.P., et al. (2013). CD106 identifies a subpopulation of mesenchymal stem cells with unique immunomodulatory properties. PLoS One 8, e59354.

Yip, S.H., Sham, P.C., and Wang, J. (2018). Evaluation of tools for highly variable gene discovery from single-cell RNA-seq data. Briefings in bioinformatics.

Yoo, K.H., Jang, I.K., Lee, M.W., Kim, H.E., Yang, M.S., Eom, Y., Lee, J.E., Kim, Y.J., Yang, S.K., Jung, H.L., et al. (2009). Comparison of immunomodulatory properties of mesenchymal stem cells derived from adult human tissues. Cell Immunol 259, 150–156.

Zhang, J., Huang, X., Wang, H., Liu, X., Zhang, T., Wang, Y., and Hu, D. (2015). The challenges and promises of allogeneic mesenchymal stem cells for use as a cell-based therapy. Stem Cell Res Ther 6, 234.

Zuk, P.A., Zhu, M., Ashjian, P., De Ugarte, D.A., Huang, J.I., Mizuno, H., Alfonso, Z.C., Fraser, J.K., Benhaim, P., and Hedrick, M.H. (2002). Human adipose tissue is a source of multipotent stem cells. Mol Biol Cell 13, 4279–4295.

