## Supplemental Information for "Single-cell RNA-seq highlights heterogeneity in human primary Wharton’s Jelly mesenchymal stem/stromal cells cultured *in vitro*"

#### Supplemental table legends

**Table S1 Top100 CD genes ranked by mean expression.**

**Table S2 CD genes ranked by percentage of cells with at least 1 UMI.**

**Table S3 Highly variable genes identified in WJMSCs**

**Table S4 DEGs among the six clusters.**

**Table S5 Marker genes for lineage differentiation potency score analysis.**

#### Supplemental figures

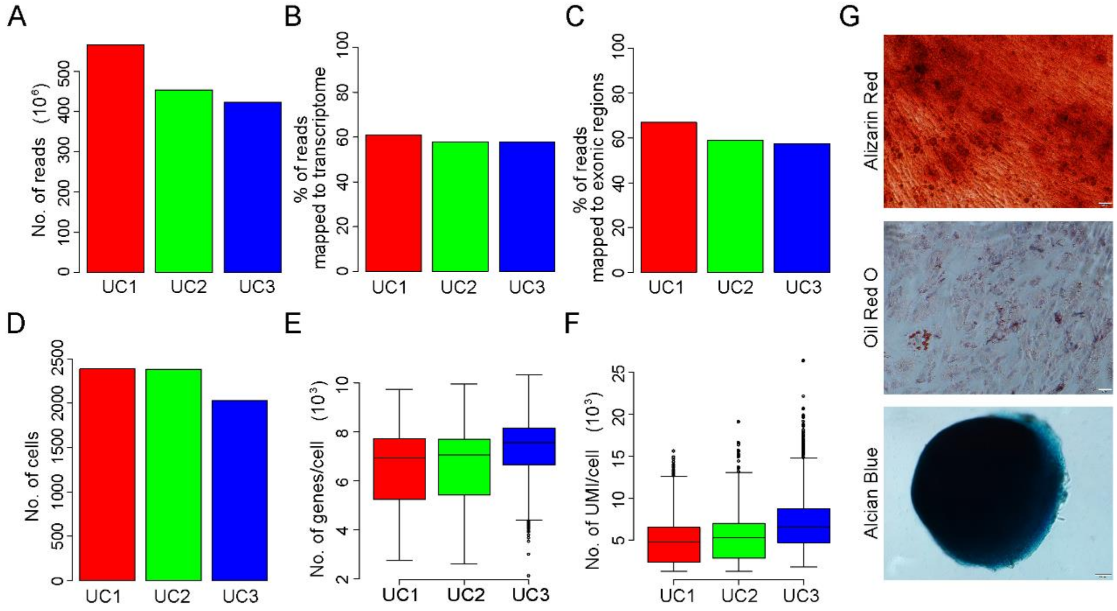


**Figure S1. Quality of the WJMSCs single-cell RNA-seq data.**

(A) Number of reads were sequenced for each of the three samples. Percentage of reads mapped to exonic (B) and mapped to transcriptome (C) for each of the three samples. (D) Number of cells were abtained for each of the three samples. Boxplot showing number of expressed genes per cell (E) and number of UMI per cell (F) for each of the three samples. (G) Tri-lineage differentiation potency of primary cultured WJ-MSCs used for scRNa-seq. Alizarin Red, Oil Red O, and Alcian Blue staining indicated the osteogenic, adipogenic, and chondrogenic differentiation, respectively.

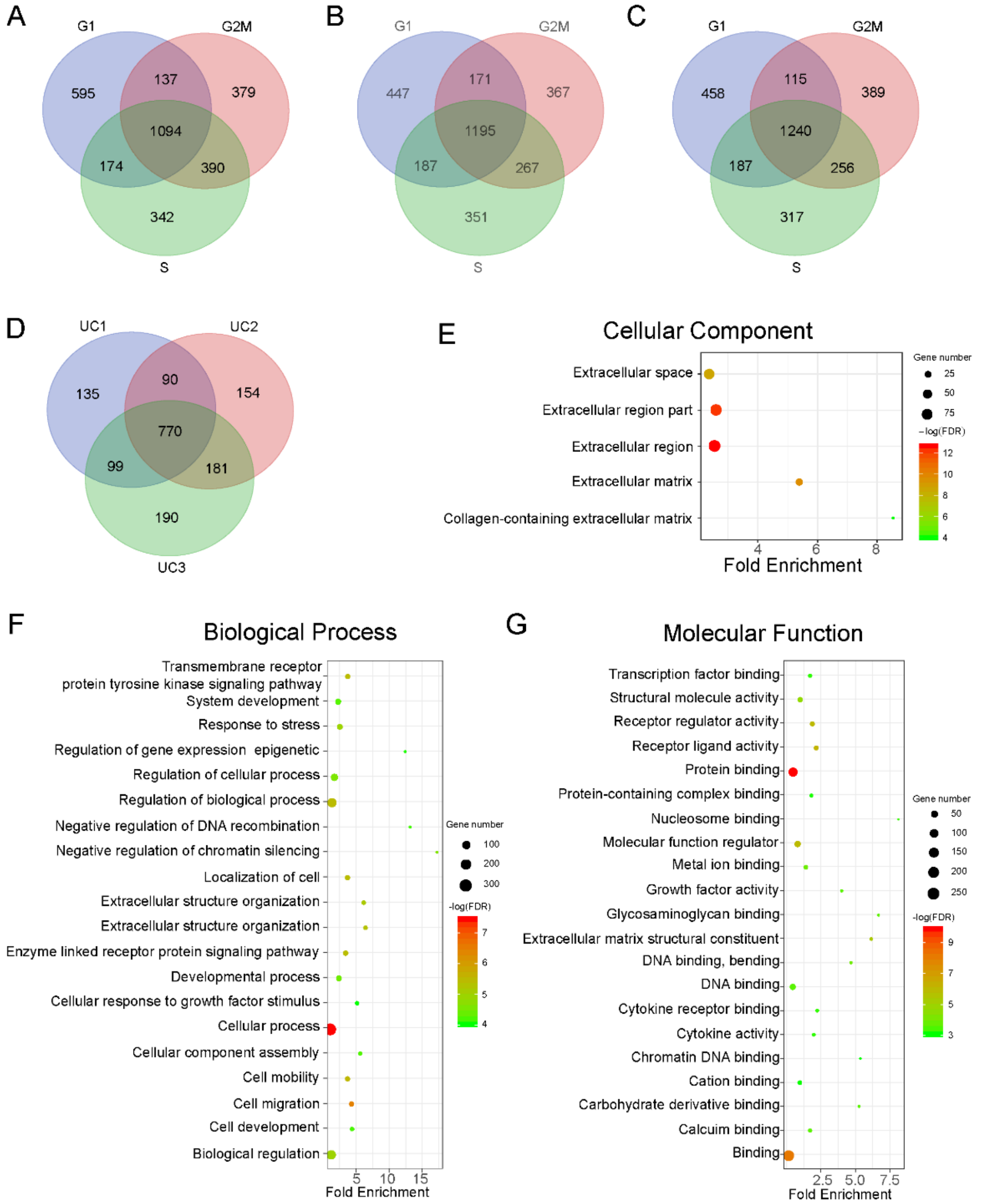


**Figure S2. Highly variable genes identification in WJ-MSCs and GO enrichment analysis.** (A) Venn diagram showing overlap of top 2000 highly variable genes among different phases for sample UC1. (B) Venn diagram showing overlap of top 2000 highly variable genes among different phases for sample UC2. (C) Venn diagram showing overlap of top 2000 highly variable genes among different phases for sample UC3. (D) Venn diagram showing overlap of highly variable genes among samples. Results of GO-slim cellular component enrichment analysis (E), GO-slim biological process enrichment analysis (F), and GO-slim functional molecular enrichment analysis for highly variable genes.

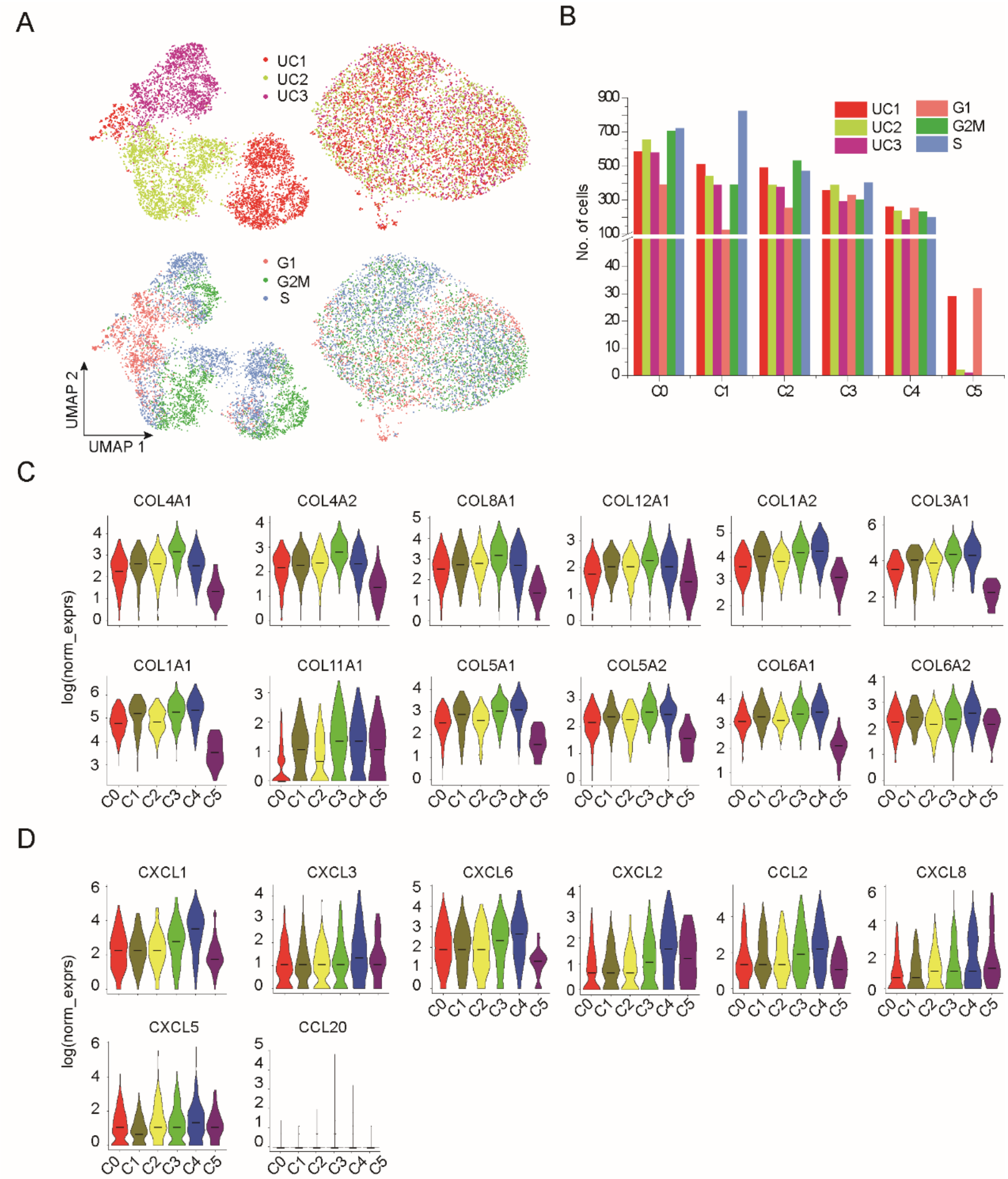


**Figure S3. Candidate subpopulations identified in WJMSCs.**

(A) UMAP showing dimension reduction before and after batch and cell cycle effect remove. left, before remove; right, after remove. (B) Histogram showing number of cells for each phase of cell cycle and sample in the candidate subpopulations. (C) Violin plots showing distribution of log normalized expression (log(norm_exprs)) values of collagen genes across the six candidate subpopulations (C0–C5). (B) Violin plots showing distribution of log(norm_exprs) values of chemokines genes across the six candidate subpopulations (C0–C5).

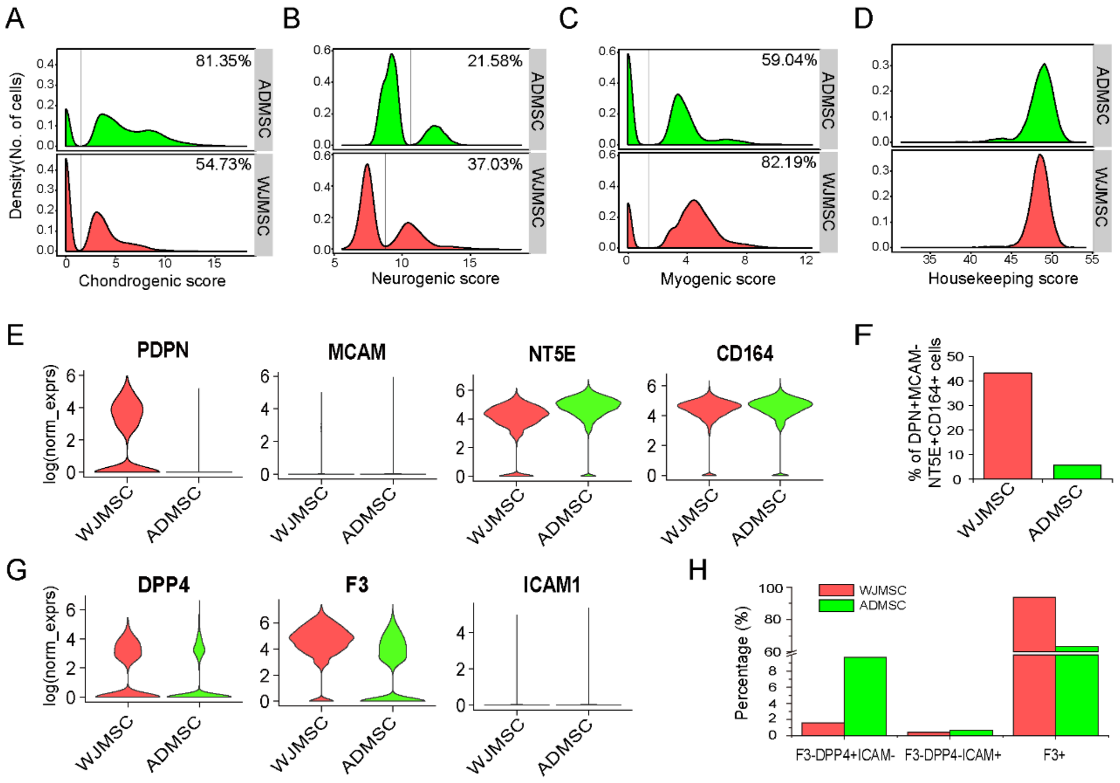


**Figure S4. Differentiation potency compared between ADMSCs and WJMSCs.**

Density distribution showing chondrogenic score(A), neurogenic score (B), myogenic score(C), and housekeeping score (D) between ADMSCs and WJMSCs. Percentage indicating proportion of cells assigned to the right side of the line. (E) Violin plot showing marker genes of SSC expressed in ADMSC and WJMSC. (F) Percentage of cells expressed SSC marker genes in ADMSC and WJMSC. (G)Violin plot showing marker genes of adipose progenitors expressed in ADMSC and WJMSC. (H) Percentage of cell expressed marker genes of adipose progenitors in ADMSC and WJMSC. F3-DPP4+ICAM-indicating phenotype of interstitial progenitor cells, F3-DPP4+ICAM- indicating phenotype of committed preadipocytes.

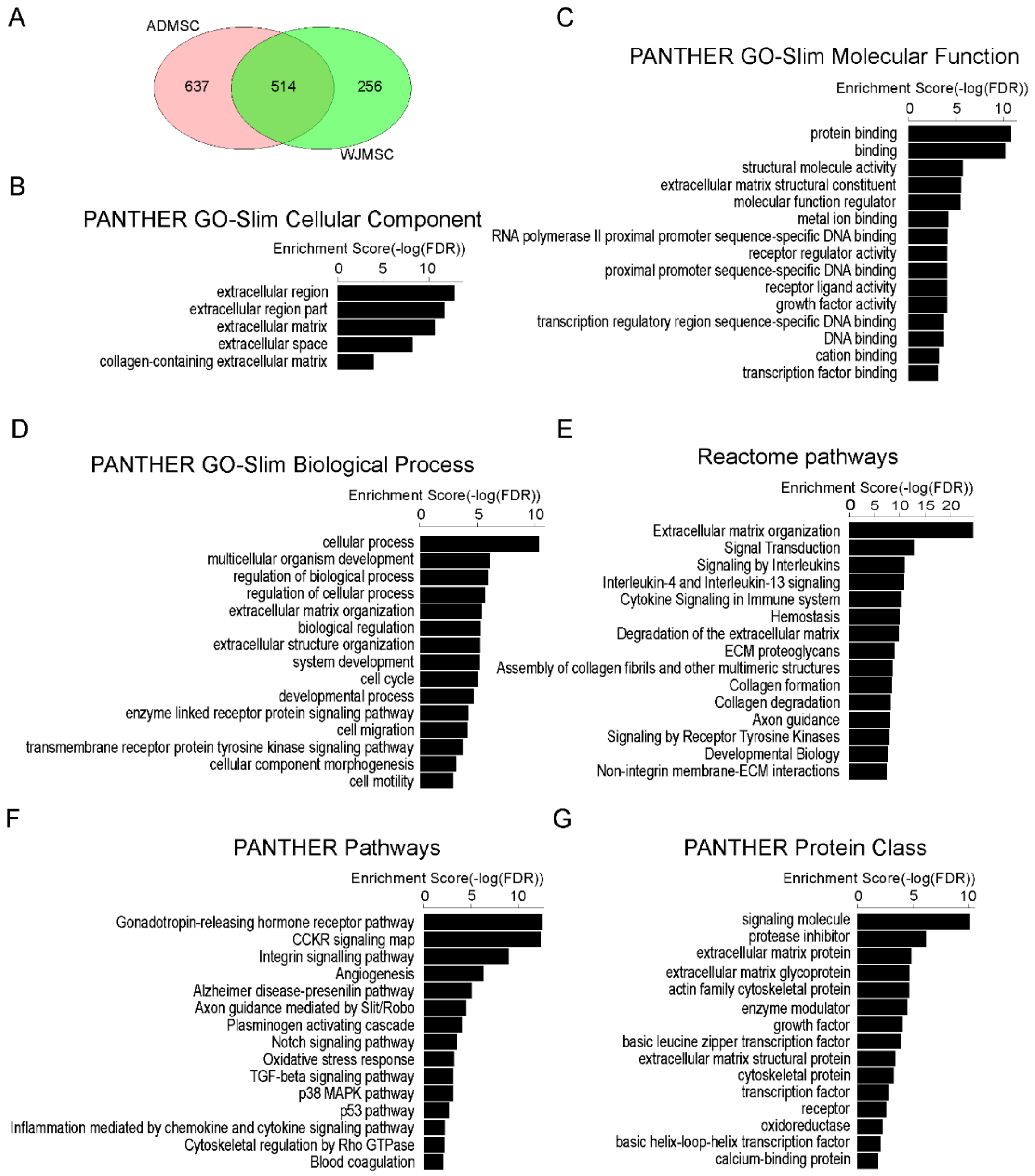


**Fig S5. Functional enrichment of highly variable genes identified in ADMSC.**

(A) Venn diagram showing overlap of HVGs between ADMSC and WJMSC. Barplots showing results of GO-slim cellular component (B), GO-slim molecular function (C), Go-slim biological process (D), Reactome pathways (E), Pathways (F), and protein class (G) enrichment analysis for HVGs identified in ADMSC. The top 15 terms ranked by -log (FDR) presented here.

### Supplemental Experimental Procedures

#### Quality Control

The scRNA-seq data was processed using cellranger-2.0.0 (https://support.10xgenomics.com) for each sample with default parameters mapping to the human GRCh38 genome expect the number of recovered cells (--expect-cells option) was set to 2 000. The resulting gene-cell UMI count matrices for each sample were then concatenated into one matrix using the “cellranger aggr” pipeline without normalization.

We filtered outlier cells using the median absolute deviation from the median total library size (logarithmic scale), total gene numbers (logarithmic scale), as well as mitochondrial percentage, as implemented in scran, using a cutoff of 3 (isOutlier, nmads = 3) (Lun et al., 2016). In total, 702 cells were removed and 6 176 high quality single cells were passed on to downstream analysis. We also filtered gene expression profile of each cell by mean-based filter. Any gene expressed across all the cells by average UMI less than 0.1 was removed, and 11 458 genes were kept for downstream analysis. Then, clean gene-cell UMI count matrix was loaded as Seurat object (Macosko et al., 2015) to manage our dataset for the further analysis. Briefly, UMI counts were used to create Seurat object by CreateSeuratObject function from R package Seurat 3.0 followed by natural log normalization using NormalizeData function with scale.factor parameter set to 1 000 000 as normalized data for genes expression comparison and figures presentation.

SRA files of public available ADMSCs (SRA:SRP148833) scRNA-seq data (Liu et al., 2019) were downloaded from https://www.ncbi.nlm.nih.gov/sra. Then SRA files were converted to fastq format and analyzed by the same pipeline used for WJMSC single cell transcriptome data.

#### Removal of Cell Cycle Effect

Considering the effect of cell cycle on genes expression, we assigned the cell cycle scores (i.e., G2/M scores and S scores) and phases (i.e. G1, G2/M, and S) for each cell on the basis of scores using function CellCycleScoring from R package Seurat based on the expression levels of a panel of phase-specific marker genes (Nestorowa and Hamey, 2016) . To remove the cell cycle effect, the S scores and G2/M scores were used to regress out cell cycle effect.

#### Highly Variable Genes Identification

Taking cell cycle and batch effects on gene expression into consideration, we created sub-datasets of cell each cycle phase for each sample, i.e. 9 sub-datasets from 3 cycle phase of 3 samples, and then performed each sub-dataset respectively for HVGs selection. We used FindVariableFeatures function in Seurat v3 (method="vst") to compute the variance of standardized values across all cells for each gene. This variance represents a measure of single cell dispersion after controlling for mean expression and can be used directly to rank the genes (Pijuan Sala et al., 2019). Here, we selected the 2 000 genes with the highest standardized variance as "highly variable" for each sub-dataset, and then chose the overlapped genes among the 9 sub-datasets to define HVGs for the following functional enrichment and dimensional reduction analysis.

#### Linear and Nonlinear Dimension Reduction

Before dimension reduction, We firstly performed normalization and variance stabilization of our data using regularized negative binomial regression (Hafemeister and Satija, 2019) with SCTransform in Seurat v3 of argument vars.to.regress to remove confounding sources of variation (variables to regress out including mitochondrial mapping percentage, number of UMI, and cell cycle scores) and argument batch_var to regress out batch effect. Then we used the scaled data of the SCT added as a new assay for the Seurat object to perform dimension reduction.

Here, we applied principal component analysis (PCA) and uniform manifold approximation and projection (UMAP) (McInnes et al., 2018) for linear and non-linear dimension reduction respectively. PCA was performed by RunPCA function in Seurat library with default parameters, except using HVGs as features augment. We used UMAP to visualize and explore data in two-dimensional coordinates, generated by RunUMAP function in Seurat. The first 20 PCs were used as input.

**Clustering the Cells**

A graph-based clustering approach (Macosko et al., 2015) were used to cluster the cells into candidate subpopulations. The first 30 PCs in the data were applied to construct an SNN matrix using the FindNeighbors function in Seurat v3 with k.param set to 20. We then identified clusters using the FindClusters command with the resolution parameter set to 0.6.

**Differential Expression Analysis**

To find differential expressed genes (DEGs) for each subpopulation, Wilcoxon Rank Sum test were performed for significant test using Seurat function FindAllMarkers for every cluster compared to all remaining cells and FindMarkers for distinguishing each other. Genes with average natural log fold change more than 0.2 and FDR less than 0.01 were assigned as DEGs.

**Tri-lineage Differentiation**

For osteogenic differentiation, MSCs were seeded into a 24-well plate at the density of 5 × 103 cells/well. When cells reached 70 % confluency, the medium was replaced with osteogenic differentiation medium (MEM-alpha medium (Thermo Fisher Scientific)，10 % FBS (Hyclone), 1 % Pen-Strep (Thermo Fisher Scientific), 100 nM dexamethasone (Sigma), 10 mM sodium β-glycero phosphate (Sigma), 0.05 mM ascorbic acid (Sigma)) and kept for 3 weeks. To assess osteogenic differentiation, Alizarin Red S staining (Sigma) was performed for the calcium-rich extracellular matrix.

For adipogenic differentiation, 1 × 104 cells were seeded per well. The cells at confluence were then treated with adipogenic differentiation medium (high glucose DMEM (Thermo Fisher Scientific), 10 % FBS (Hyclone), 0.5 mM IBMX (Sigma), 1μM dexamethasone (Sigma), 1.7 μM insulin (Sigma), 0.2 mM indomethacin (Sigma)) for 3 weeks. The cells were fixed with 4% formaldehyde solution and lipid droplets of the resultant differentiated cells were detected using Oil Red O staining (Sigma).

For chondrogenic differentiation, StemPro™ Chondrogenesis Differentiation Kit were used and performed according to the manufacturer protocol. Briefly, 1.6× 107 cells resuspended in MSC medium. Generate micromass cultures by seeding 5-μL droplets of cell solution in the center of 96-well plate wells. After cultivating micromass cultures for 2 hours under high humidity conditions, add warmed chondrogenesis media to culture vessels and incubate in 37 °C incubator with 5 % $CO_2$. After 3 weeks, fix cells with 4 % formaldehyde solution and stained with Alcian Blue (Sigma).

#### Supplemental Reference

Hafemeister, C., and Satija, R. (2019). Normalization and variance stabilization of single-cell RNA-seq data using regularized negative binomial regression. bioRxiv, 576827.

Liu, X., Xiang, Q., Xu, F., Huang, J., Yu, N., Zhang, Q., Long, X., and Zhou, Z. (2019). Single-cell RNA-seq of cultured human adipose-derived mesenchymal stem cells. Scientific data 6, 190031.

Lun, A.T., McCarthy, D.J., and Marioni, J.C. (2016). A step-by-step workflow for low-level analysis of single-cell RNA-seq data with Bioconductor. F1000Research 5, 2122.

Macosko, E.Z., Basu, A., Satija, R., Nemesh, J., Shekhar, K., Goldman, M., Tirosh, I., Bialas, A.R., Kamitaki, N., Martersteck, E.M., et al. (2015). Highly Parallel Genome-wide Expression Profiling of Individual Cells Using Nanoliter Droplets. Cell 161, 1202-1214.

McInnes, L., Healy, J., and Melville, J. (2018). Umap: Uniform manifold approximation and projection for dimension reduction. arXiv preprint arXiv:180203426.

Nestorowa, S., and Hamey, F.K. (2016). A single-cell resolution map of mouse hematopoietic stem and progenitor cell differentiation. 128, e20-31.

Pijuan Sala, B., Diamanti, E., Shepherd, M., Laurenti, E., Wilson, N.K., Kent, D.G., Gottgens, B., Stuart, T., Butler, A., Hoffman, P., et al. (2019). Comprehensive Integration of Single-Cell Data. Blood 177, 1888-1902.e1821.
